# Improving RNA branching predictions: advances and limitations

**DOI:** 10.1101/2021.02.04.429782

**Authors:** Svetlana Poznanovíc, Carson Wood, Michael Cloer, Christine Heitsch

**Affiliations:** School of Mathematical and Statistical Sciences, Clemson University, Clemson, SC 29634, USA; School of Mathematics, Georgia Institute of Technology, Atlanta, GA 30308, USA

**Keywords:** secondary structure, NNTM, multiloops, branching parameters

## Abstract

Minimum free energy prediction of RNA secondary structures is based on the Nearest Neighbor Thermodynamics Model. While such predictions are typically good, the accuracy can vary widely even for short sequences, and the branching thermodynamics are an important factor in this variance. Recently, the simplest model for multiloop energetics — a linear function of the number of branches and unpaired nucleotides — was found to be the best. Subsequently, a parametric analysis demonstrated that per family accuracy can be improved by changing the weightings in this linear function. However, the extent of improvement was not known due to the ad hoc method used to find the new parameters. Here we develop a branch-and-bound algorithm that finds the set of optimal parameters with the highest average accuracy for a given set of sequences. Our analysis shows that the previous ad hoc parameters are nearly optimal for tRNA and 5S rRNA sequences on both training and testing sets. Moreover, cross-family improvement is possible but more difficult because competing parameter regions favor different families. The results also indicate that restricting the unpaired nucleotide penalty to small values is warranted. This reduction makes analyzing longer sequences using the present techniques more feasible.

## 1. Introduction

Accurate prediction of RNA base pairings from sequence remains a fundamental problem in bioinformatics. While new methods continue to advance the ribonomics research frontier [1–9], experimentalists still obtain useful functional insights from the classical minimum free energy (MFE) secondary structure predictions [10].

An RNA secondary structure is a set of intra-sequence base pairs. For thermodynamic prediction purposes, the target pairings are typically canonical, i.e. Watson-Crick or wobble, and pseudoknot-free. (Removing these constraints, especially the latter, are active areas of research in the field.) Given a secondary structure, its free energy change from the unfolded sequence can be approximated under the Nearest Neighbor Thermodynamic Model (NNTM); this model and the evolution of its many parameters are cataloged in an online database [11]. Under the NNTM, when given a sequence, an MFE secondary structure can be computed efficiently using dynamic programming [12–15].

Historically, the prediction accuracy is high on average for sequences of length 700 nucleotides or less [16]. In particular, it was found that an average (with standard deviation) of 83.0% (±22.2) of pairings in 484 transfer RNA (tRNA) secondary structures, totaling 10,018 base pairs (bp) and 37,502 nucleotides (nt), were predicted correctly. Likewise, 77.7% (±23.1) of pairings in 309 5S ribosomal RNA (rRNA) secondary structures, totaling 10,188 bp and 26,925 nt, were predicted correctly. While this clearly supports the value of MFE predictions, it also highlights that even at this scale of sequence lengths, i.e. ∼76 nt for tRNA and ∼120 for 5S rRNA, the prediction accuracy for an individual sequence can be low.

Recent results have demonstrated that it is possible to obtain a statistically significant increase in MFE prediction accuracy on a diverse training set of 50 tRNA sequences and 50 5S rRNA by changing the thermodynamic cost of branching [17]. Recall that an RNA secondary structure is composed of different substructures, which are scored by different components of the NNTM objective function. The substructures known as multiloops (or branching junctions) have three or more helices which radiate out as branches. A tRNA molecule has one central multiloop with four branches, while 5S rRNA has one with three. Although these branching loops are a critical aspect of the overall molecular conformation, they remain one of the most difficult aspects to predict accurately [18,19].

Previously, mathematical techniques from discrete optimization and geometric combinatorics were used to completely characterize all the secondary structures which were optimal for all possible combinations of different branching parameters on the chosen training sets [17]. It was found that 89% of tRNA and 90% of 5S rRNA predictions could be improved by altering the branching parameters from the default values. (Those which did not improve already had an accuracy well above average.) Critically, though, achieving this improvement *simultaneously*, i.e. for the same set of new parameters, is not possible; the intersection of all the “best possible” set of parameters for each sequence is empty.

At the time, an ad hoc combinatorial method was used to identify large combinations of non-empty intersections among the individual “best possible” sets for a given collection of training sequences. It was demonstrated that the branching parameters obtained in this way yielded a statistically significant improvement in MFE prediction accuracy over the existing values on tRNA, on 5S rRNA, and on the total 100 sequence training set, respectively. However, a significant gap remained between the average (known to be unobtainable) of the maximum attainable individual accuracies with modified branching parameters, and the best ad hoc values found.

The new method and associated results presented here eliminate that gap by giving a branch-and-bound algorithm, and an effective implementation, for finding parameters with the optimal MFE prediction accuracy across the given collection of RNA sequences. Somewhat surprisingly, we find that the improvement in prediction accuracy between the new branch-and-bound (BB) parameters and previous ad hoc (AH) ones is *not* statistically significant. To test this conclusion, we computed the prediction accuracies for the different branching parameters under consideration on a much larger set of 557 tRNA and 1283 5S rRNA sequences [20] from the Mathews Lab (U Rochester). We again saw no significant difference between the AH and BB parameters in MFE prediction accuracy for the testing sequences.

Moreover, we also confirmed that the new parameters give a statistically significant improvement over the current ones on the type of testing sequences, i.e. tRNA, 5S rRNA, or both, for which the parameters were trained. In conjunction, these results suggest that there may be a relatively large set of branching parameters which yield equivalent prediction accuracies that improve over the current ones. To move forward in identifying the scope of these parameters, we confirm that the current empirical strategy of focusing on the trade-off between two of the three branching parameters is well-substantiated by our analysis.

## 2. Materials and Methods

We briefly sketch some background in the parametric analysis of RNA branching relevant to the current work before giving the new branch-and-bound algorithm. We conclude with information about the training and testing sequences used.

### 2.1 Parametric analysis of RNA branching

An RNA secondary structure decomposes into well-defined substructures. Our focus here is on the ones known as multiloops or branching junctions. Such a substructure has 3 or more helical arms which radiate out as branches. The classical tRNA cloverleaf has a single multiloop with four branches, while the single 5S rRNA one has only three. A parametric analysis seeks to understand how the MFE prediction depends on the parameters used in the thermodynamic optimization. Here we focus on the three which govern the entropic cost of loop branching.

#### 2.1.1 Branching parameters

We use the term *branching parameters* to refer to the three (learnt) parameters (*a, b, c*) in the initiation term in the multiloop scoring function:

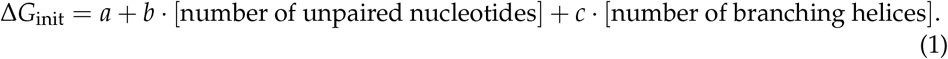

The initiation term, together with the “stacking” energies of adjacent single-stranded nucleotides on base pairs in the loop, is used to approximate the multiloop stability under the NNTM. The stacking energies are based on experimental measurements [16, 21], but the linear form of the initiation term was originally chosen for computation efficiency [21]. This simple entropy approximation has been shown to outperform other more complicated models for multiloop scoring in MFE prediction accuracy [22]. In this we work we focus on possible improvements of the MFE prediction by changing the branching parameters.

#### 2.1.2 Standard branching parameters

The NNTM has evolved over time, and the loop initiation parameters have changed with each major revision. Here they will be denoted T89 [21], T99 [16], and T04 [22] as in Table 2. On the two families considered, the T99 values were the most accurate overall, so they will be the primary point of comparison for the results. The older and newer values are also listed, and some trade-offs among the three will be addressed in the discussion.

**Table 2.** Parameter values and MFE prediction for the 50 tRNA training sequences and the 50 5S rRNA, as well as over both families.

|  | Parameters |  |  | tRNA |  | 5S rRNA |  | Both |  |
| --- | --- | --- | --- | --- | --- | --- | --- | --- | --- |
|  | a | b | c | avg | std | avg | std | avg | std |
| T89 | 4.6 | 0.4 | 0.1 | 0.41 | 0.25 | 0.69 | 0.24 | 0.55 | 0.28 |
| T99 | 3.4 | 0 | 0.4 | 0.52 | 0.30 | 0.63 | 0.24 | 0.58 | 0.27 |
| T04 | 9.3 | 0 | -0.6 | 0.45 | 0.28 | 0.64 | 0.24 | 0.54 | 0.28 |
| max | n/a | n/a | n/a | 0.91 | 0.10 | 0.81 | 0.09 | 0.86 | 0.11 |
| AHt | 10.9 | -0.1 | -2.6 | 0.74 | 0.24 | 0.52 | 0.21 | 0.63 | 0.25 |
| AHs | -8.5 | 0.3 | 4.5 | 0.36 | 0.18 | 0.71 | 0.22 | 0.53 | 0.27 |
| BBt | 17 | -0.3 | -4.5 | 0.75 | 0.22 | 0.45 | 0.17 | 0.60 | 0.25 |
| BBs | -5.7 | 0.2 | 3.5 | 0.37 | 0.19 | 0.73 | 0.21 | 0.55 | 0.27 |
| AHb | 12.2 | 0.2 | -2.9 | 0.71 | 0.27 | 0.62 | 0.21 | 0.66 | 0.25 |
| BBb | 9.3 | -0.1 | -1.7 | 0.73 | 0.25 | 0.59 | 0.23 | 0.66 | 0.25 |

#### 2.1.3 Precision of branching parameters

For technical reasons, the parametric analysis is performed over the rationals with very high precision. However, all current thermodynamic optimization methods use rather low precision by comparison. In particular, the branching parameters are specified to one decimal place. For the presentation of results and discussion of their implications, the new branching parameters used were first rounded to one decimal place.

#### 2.1.4 Branching polytopes

The branching polytopes were developed in [23]. They allow for parametric analysis of the portion of the NNTM used for scoring the multiloops. We give a short background here only to explain the terms that we use in the description of the branch-and-bound search for the optimal parameters later.

To a secondary structure *S*, we associate a quadruple (*x, y, z, w*) of real numbers, called the *branching signature* [17]. The components of the signature are: *x* -the number of multiloops in *S, y* -the total number of unpaired nucleotides in those multiloops, *z* -the total number of branching helices around those multiloops, and *w* -a remainder term which includes all other components of the Δ*G* calculation for *S* under the NNTM except those involving *a, b*, and *c*. The free energy of the structure (1) can then be expressed as △*G*_*S*_ = *ax* + *by* + *cz* + *w*. The branching polytope for a given RNA sequence is the convex hull of the signatures of all possible secondary structures for *R*.

Through a duality from convex geometry, to the branching polytope we associate a polyhedral fan in 4D, called the normal fan of the branching polytope. The intersection of the normal fan with the *d* = 1 plane produces a subdivision of the 3D parameter space into convex regions. These regions are of great significance to analyzing the effects of varying the branching parameters in the NNTM. Namely, the branching parameters (*a, b, c*) from the same region yield the same MFE structures, while for parameters from different regions the model produces different predictions.

### 2.2 Branch-and-bound algorithm

Let Reg_*R*_ be the (finite) list of the regions associated to an RNA sequence *R*. The average number of regions for the training data is: 517 for tRNA and 2109 for 5S rRNA. The part of parameter space that contains the optimal parameters for a given set of sequences 𝒟 can be found by considering the intersections ∩_*R*∈D_Reg_*R*_ [*k*_*R*_], for all possible combinations of the indices *k*_*R*_. Due to size, exhaustive search for the optimal region is not feasible -we use a branch-and-bound algorithm instead. We first present the basic idea behind the algorithm and how the merging and pruning steps are performed for two sequences. Then we explain how this idea can be extended to a larger training set. The straightforward extension, however, is not efficient for 50 sequences, so we perform merges along a binary tree. We also explain how performing certain pre-processing steps which require initial overhead time significantly improve the total running time.

#### 2.2.1 Basic idea behind the algorithm

Suppose we are optimizing the parameters for two RNA sequences. In this case, we are interested in all the pairwise intersections Reg_1_[*k*_1_] ∩ Reg_2_[*k*_2_] of the regions that correspond to the two training sequences. Most of the intersections are empty and should thus be discarded. However, checking for nonempty intersections is computationally expensive. Therefore, before we compute intersections, we check whether the intersection could possibly provide an improvement in the prediction.

The inputs for the algorithm are the polytope for each sequence, the optimal structures that correspond to the vertices of each of the polytopes, and a lower bound *L* for the best average accuracy. Better lower bounds improve the running time -we used the accuracy we knew we could achieve with the ad hoc parameters. For *i* = 1, 2, let Acc_*i*_(Reg_*i*_ [*j*]) denote the prediction accuracy for sequence *i* when parameters from its *j*-th region are used. This value can be computed from the input by exhaustive search through the possible optimal structures. We order the regions in the list Reg_*i*_ by decreasing accuracy. Let *U*_*i*_ = Acc_*i*_(Reg_*i*_ [1]) be the maximal attainable accuracy for sequence *i*. The region Reg_1_[*j*] can be discarded from consideration in the search for optimal parameters unless

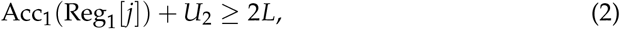

If (2) is satisfied, we say Reg_1_[*j*] passes the pruning test. Note that since the regions in Reg_1_ are listed by accuracy, once *j* is large enough so that the pruning test fails, we can conclude that it would fail for all remaining regions from Reg_1_ and we remove them from future consideration.

If Reg_1_[*j*] passes the pruning test, in the next step we consider which of its inter-sections Reg_1_[*j*] ∩Reg_2_[*k*] with regions from the second sequence are nonempty and yield accuracy better then *L*. We consider these candidate regions starting with the lowest *j* and, for a fixed *j*, we check for *k* in increasing order. The first nonempty intersection found, say Reg_1_[*j*_0_] ∩ Reg_2_[*k*_0_], is a region for which the average accuracy is (Acc_1_(Reg_1_[*j*_0_]) + Acc_2_(Reg_2_[*k*_0_]))/2, so we update our lower bound *L*. Since the regions in Reg_2_ are listed by accuracy, we start checking for the next value of *j*. Then before we compute other candidate regions Reg_1_[*j*] ∩ Reg_2_[*k*], we check whether

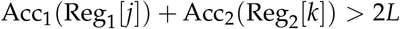

and only do polytope calculations if this inequality is satisfied. Each time we find a new nonempty intersection, the value *L* is updated as before. At the end, the value *L* is the best attainable accuracy and the last nonempty intersection that was computed is the part of parameter space that yields this accuracy.

Note that the time spent for ordering the lists Reg_1_ and Reg_2_ at the beginning saves time in performing polytope intersections later.

#### 2.2.2. Extending to *N* sequences

The regions ∩_*R*∈ 𝒟_Reg_*R*_ [*k*_*R*_] of interest can be formed by sequentially intersecting all the regions from Reg_1_ corresponding to the first sequence with all the regions from Reg_2_ corresponding to the second sequence, then taking all the nonempty pairwise intersections and intersecting them with the regions from Reg_3_, etc. Figure 1 is an illustration of the linear order in which these intersections can be formed in the simplest version of the algorithm.

**Figure 1.**
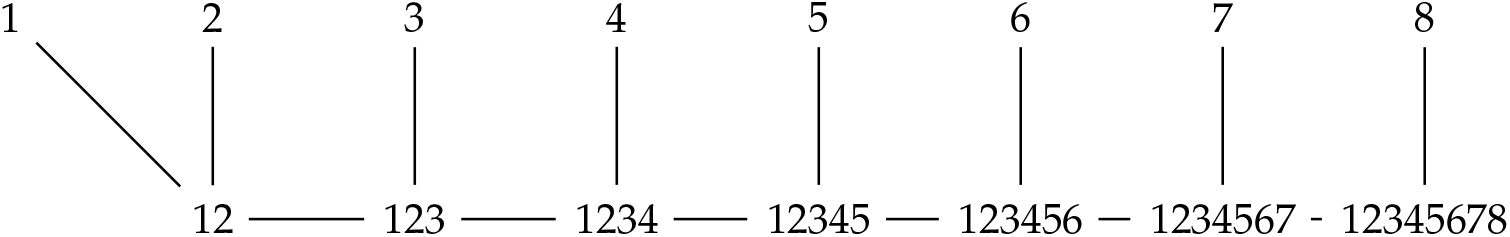
Order of merges under the basic branch-and-bound algorithm.

When checking which regions from the first sequence should be considered in forming the intersections, the criterion (2) is replaced with

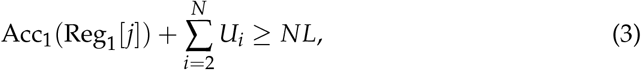

where *U*(*i*) is the maximal attainable accuracy for the *i*-th sequence, computed from the input data.

In the first merge illustrated in Figure 1, for each Reg_1_[*j*] which passes this pruning test, we consider regions Reg_2_[*k*] from the second sequence. If

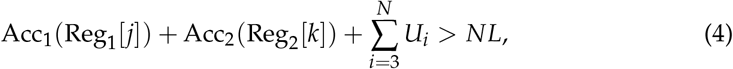

we check whether Reg_1_[*j*] ∩Reg_2_[*k*] is nonempty, and if so we consider it in the next merge, etc. In the final merge, similarly to the case of 2 sequences, we can exploit the fact that we know that the accuracy for a nonempty intersection 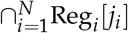 is 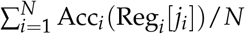, so each time we find a nonempty intersection, we update the value of *L* and use this new value to prune some of the remaining candidates before we actually perform polytope calculations.

In fact, the polytope calculations being the most time consuming part of the process, it turns out that it pays off to invest time in finding better upper bounds to replace the *U*_*i*_’s in the pruning steps. Therefore, before we start the merges, we compute the maximal attainable accuracy for sequence *i* for each region Reg_1_[*j*], denoted by Max_*i*_(Reg_1_[*j*]). In order to do this, we identify which intersections Reg_1_[*j*] ∩ Reg_*i*_ [*k*] are nonempty and take the maximal Acc_*i*_(Reg_*i*_ [*k*]). The pruning test (3) is then replaced by

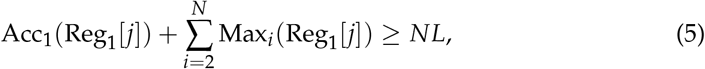

while (4) is replaced by

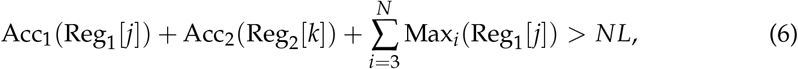

etc.

#### 2.2.3. Binary merge

When merging regions for *k* tRNA sequences, we need to consider intersections of *k* polyhedral regions, so it is important that we maximize the number of those that can get pruned before any polytope calculations are performed. Therefore, in our implementation we replace the linear merge from Figure 1 by a binary merge order, illustrated in Figure 2. Each node of the tree represents a step in which intersections of regions from two sets of sequences are being considered.

**Figure 2.**
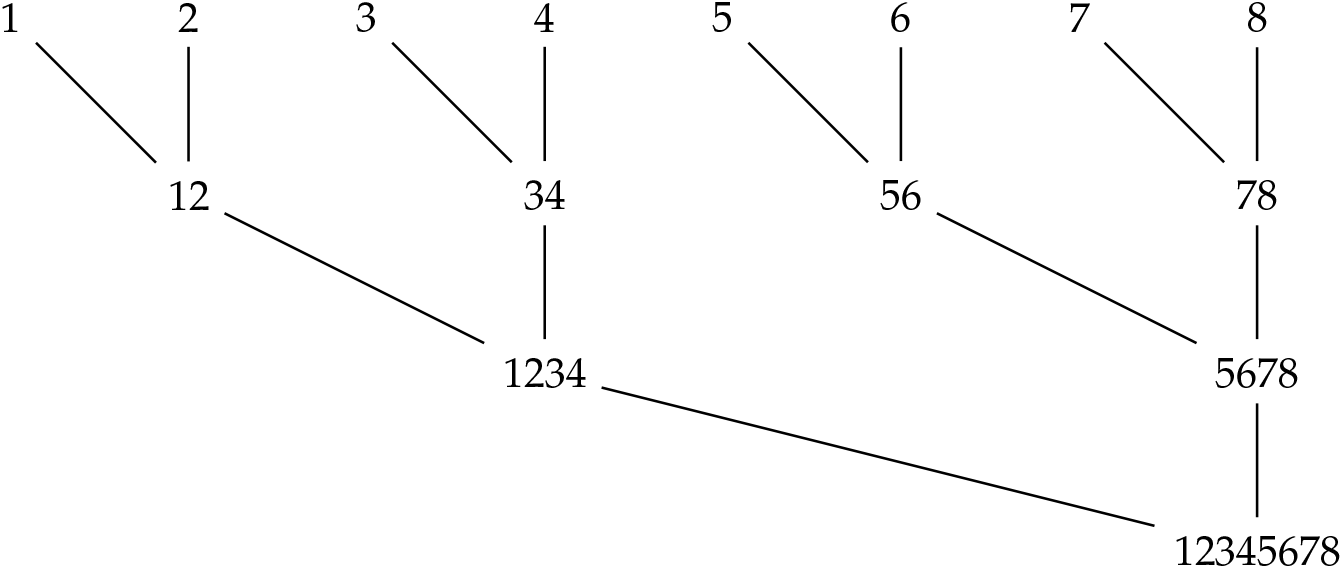
Improved merging order.

The improved upper bounds that we compute at the beginning in order to have more efficient pruning are Max_*k*_(Reg_*i*_ [*j*]) -the maximal accuracy for sequence *k* under parameters from the region Reg_*i*_ [*j*]). While it may seem like each of these values requires Reg_*k*_ polytope intersections, here we exploit the fact that the regions |Reg_*k*_| are ordered by accuracy, thereby significantly reducing the computing time. Namely, we consider intersections Reg_*k*_ [*l*] ∩ Reg_*i*_ [*j*]) ordered by increasing *l* and when the first nonempty intersection is found, we can take Max_*k*_(Reg_*i*_ [*j*]) = Acc_*k*_(Reg_*k*_ [*l*]).

In fact, the values Max_*i*_(Reg_*k*_ [*j*]) computed can be used before merges start to reduce the number of regions we consider for each sequence ^1^. This is another instance where we pay slight overhead at the beginning in order to save time on calculating polytope intersections later. Namely, in a pre-processing step at the beginning, right after the values Max_*i*_(Reg_*k*_ [*j*]) are calculated, region Reg_*i*_ [*j*] is discarded from future consideration unless

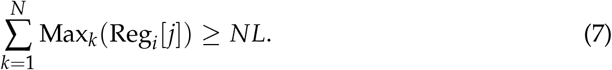

During merges, when the regions from sequences *i* ∈ *I* are being merged, the intersection ∩_*i*∈*I*_Reg_*i*_ [*k*_*i*_] is not computed unless it passes the pruning test

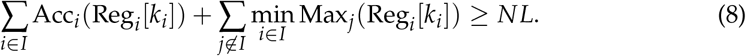

If the intersection passes the test, then we check that it is nonempty before we save it to be considered in future merges. However, when training on 100 sequences this still leads to checking a prohibitively large number of intersections of a large number of polytopes. For that reason, when the number of sequences being merged is at least ten, we use an additional test before we verify that ∩_*i*∈*I*_Reg_*i*_ [*k*_*i*_] is nonempty. Namely, suppose the merge *I* comes from two branches, *I* = *I*_1_ ∪ *I*_2_. Then, by construction, 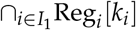 and 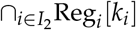 are both nonempty. However, 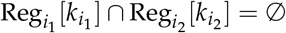 implies that the whole intersection is empty, even though the opposite is not true. Therefore, we verify which pairwise intersections 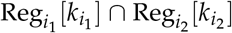, for *i*_1_ ∈ *I*_1_, *i*_2_ ∈ *I*_2_, are nonempty for all the regions 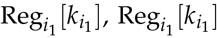 that appear in the two lists of intersections for the sequences from the sets *I*_1_ and *I*_2_, respectively, that are being merged at this step. We then check whether ∩_*i*_∈_*I*_Reg_*i*_ [*k*_*i*_] can be declared empty based on this information, and if not, then we compute the intersection. Most pairwise intersections considered here are empty, and when |*I*|≥ 10, the number of regions to be considered is small enough that the overhead time used to compute the pairwise intersections at the beginning of the merge is a good trade-off for the time saved in checking the nonempty intersections when the pruning test is passed.

### 2.3. Data analysis

Parameters obtained from the new branch-and-bound algorithm were trained on sequences from the previous study and tested on a much larger set available from the Mathews Lab (U Rochester) to determine the extent of possible improvement in MFE prediction accuracy by modifying the branching parameters.

#### 2.3.1. Accuracy measure

To determine the accuracy of a prediction we compare with a pseudoknot-free native secondary structure *S* from which the noncannonical base pairings have been excluded. For an MFE prediction *S′* for that RNA sequence, we score the accuracy as the *F*_1_-measure:

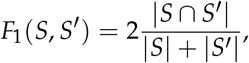

where |*S*| and |*S′*|are the number of base pairs in *S* and *S′*, respectively, and |*S* ∩*S′*| is the number of true positive base pairs common to both structures. The minimum value 0 means no accurately predicted base pairs, while 1 means perfect prediction. The accuracy of a multiloop initiation parameter triple for a sequence is the average over all possible MFE secondary structures for that fixed (*a, b, c*).

#### 2.3.2. Testing and training sequences

Two sets of sequences were used in this study. The first set was used for training, and consisted of the 50 tRNA and 50 5S rRNA sequences from the previous study [17]. (A complete list with Accession number is given in the Supplementary data of the previous paper.) These sequences were obtained from the Comparative RNA Web (CRW) Site [24]. By design, they were chosen so that their MFE prediction accuracies and GC content are as uniformly distributed as reasonably possible. The GC content was used to ensure that sequences with very similar MFE accuracies were sufficiently different to generate a diverse training set.

The second set was used for testing, and consisted of the 557 tRNA and 1283 5S rRNA sequences from a larger benchmarking set first used in [20,22], which supersedes previous ones [16]. Summary statistics for some characteristics like sequence length, MFE accuracy under the Turner99 branching parameters, GC content, and number of native base pairs (bp) for both data sets broken down by family are given in Table 1. While the differences in sequence length between the training and testing data were significant for both families (tRNA: *p* = 0.0003, 5S rRNA: *p* = 0.0000), the differences in MFE accuracy were not (tRNA: *p* = 0.0846, 5S rRNA: *p* = 0.2659). Differences in both GC content and number of base pairs were mixed; the former had *p* = 0.0018 for tRNA but *p* = 0.0883 for 5S rRNA, whereas the latter was *p* = 0.2707 and *p* = 0.0000. As will be seen in the results section, the training data seemed to represent well the testing sequences despite any differences in the composition of the sets.

**Table 1.** Per family characteristics of the training and testing data sets.

| Type | Family | Num | Seq length |  | MFE accuracy |  | GC content |  | Native bp |  |
| --- | --- | --- | --- | --- | --- | --- | --- | --- | --- | --- |
|  |  |  | avg | std | avg | std | avg | std | avg | std |
| Training | tRNA | 50 | 74.38 | 1.89 | 0.52 | 0.30 | 0.59 | 0.08 | 20.36 | 1.21 |
| Training | 5S rRNA | 50 | 121.38 | 3.62 | 0.63 | 0.24 | 0.58 | 0.06 | 35.18 | 3.04 |
| Testing | tRNA | 557 | 77.10 | 5.21 | 0.58 | 0.24 | 0.54 | 0.11 | 20.55 | 1.15 |
| Testing | 5S rRNA | 1283 | 118.71 | 3.49 | 0.59 | 0.24 | 0.57 | 0.05 | 33.61 | 2.38 |

## 3. Results

We first address the extent of improvement possible when the branching parameters are trained on a specific family, either tRNA or 5S rRNA. We then consider the improvement possible across both training families simultaneously. The training results are tested against a much larger set of 1840 sequences, and the conclusions are found to hold. Importantly, the demonstrates that the method is not overfit to the training data. Finally, the relevance of parameter precision to the results is addressed.

### 3.1. Improving per family prediction accuracy

Previously [17], mathematical methods were used to find the best of all possible combination of branching parameters (*a, b, c*) for each individual training sequence. Averaging these per sequence accuracies over their family, or the whole training set, yields the “max” accuracy listed in Table 2. These numbers are an upper bound on the improvement possible by modifying the branching parameters for MFE prediction accuracy on these training (sub)sets. We note that the upper bound is known not to be achievable, since the common intersection (even within a family) of the best per sequence parameter values is empty.

A lower bound on the improvement was established [17] by ad hoc means, that is by identifying large sets of sequences from each family which did have such a common intersection. Seven such large sets were found for tRNA and four for 5S rRNA. These combinations of possible parameters were considered for each family, and the one with best average accuracy over the whole family was reported. We refer to these parameter combinations here as AHt and AHs, for the tRNA and 5S rRNA families respectively. It was found that the AHt and AHs accuracy on their respective family, listed in Table 2, was a statistically significant improvement over the T99 parameters.

However, there was a sizable gap between the upper and lower bounds for the potential accuracy improvements for each family considered. The branch-and-bound algorithm presented here was implemented to determine where in this range the best possible per family accuracy lay. The new parameters are listed in Table 2 as BBt and BBs along with the corresponding accuracies.

As a technical aside, the best possible (*a, b, c*) region for 5S is unbounded in the (−3, 0, 1) direction. The BBs parameters reported are the centroid of the finite 2D face. We also considered a strictly interior point BBs+(− 3, 0, 1) = (−8.7, 0.2, 4.5), which gave nearly identical accuracy for tRNA. Interestingly, this new point is very close to AHs, with a distance of (0.2, 0.1, 0).

The differences between the ad hoc and branch-and-bound parameter accuracies within the family on which the parameters were optimized are not significant, with *p* = 0.7699 for tRNA and *p* = 0.7021 for 5S rRNA. In other words, the maximum possible accuracy *per family* over the training set sequences is essentially the best ad hoc accuracy, and far from the average over the per sequence maximums. This closes the gap from the previous analysis, while opening the door to new questions as discussed in the next section.

### 3.2. Improving cross family prediction accuracy

It was observed before that the accuracy of the new AHt and AHs parameters on the other family is clearly worse than the Turner parameters. Consequently, a “best both” combination, denoted here AHb, was also identified from among the 11 possible ad hoc ones considered, 7 for tRNA and 4 for 5S rRNA. The improvement over the T99 parameters on the whole training set accuracy is significant (*p* = 0.0200). The AHb parameters raise the tRNA accuracy by an amount statistically indistinguishable (*p* = 0.5243) from AHt, while the AHb accuracy for the 5S rRNA training sequences was indistinguishable (*p* = 0.7308) from T99.

To address the problem of improving the prediction accuracy simultaneously for both families, the branch-and-bound algorithm was run on the full 100 sequence training set. The parallelization is implemented in such a way that three combinations of “near optimal” parameters were detected, along with the absolute best one. When the parameters are rounded to 1 decimal precision, 3 of the 4 combinations give identical 0.66 average accuracy when rounded to 2 decimal places, and the fourth is only 0.02 less. Given this, we report as the BBb parameters the combination with 0.66 average accuracy which was most dissimilar to AHb. The other 2 combinations were AHb +(− 0.3, 0, 0.1) and +(−0.1, 0, 0), although AHb was originally optimized for tRNA and *not* the full training set. The fourth was more similar to BBb but the combination (9.7, − 0.2, − 1.8) was noticeably worse for 5S by 0.05 with only a 0.01 improvement for tRNA.

These results illustrate that, while it is possible to improve over the current T99 prediction accuracy by a statistically significant amount, doing so simultaneously over two different families is more challenging than improving the per family accuracy. Additionally, the best possible accuracy over the full training set is the same as the previous ad hoc one.

### 3.3. Results for testing data

We note that the summary statistics for “Both” families in Table 3 were computed as a weighted average and standard deviation, so that each family contributed 50% to the statistic although there were more than twice as many 5S rRNA as tRNA sequences in the testing set. With a total of 1840 sequences, the difference in accuracy over the whole data set for the T99 parameters versus the AHb or BBb is significant; the Tukey HSD Post-hoc Test following an ANOVA were both *p* = 0.0000 while the differences between AHb and BBb were not (*p* = 0.5957). The AHb and BBb parameters performed just as well on the tRNA testing sequences as on the training ones (ANOVA *p* = 0.9008), although the 5S accuracy was less good (ANOVA *p* = 0.0051). A Tukey HSD Post-hoc Test found significant differences between the AHb training accuracy and the testing one (*p* = 0.0420) as well as the BBb testing one (*p* = 0.0156) for the 5S family, but the other four pairwise comparisons were indistinguishable. We also note that the parameters trained on 5S sequences have lower accuracy on the testing data.

**Table 3.**
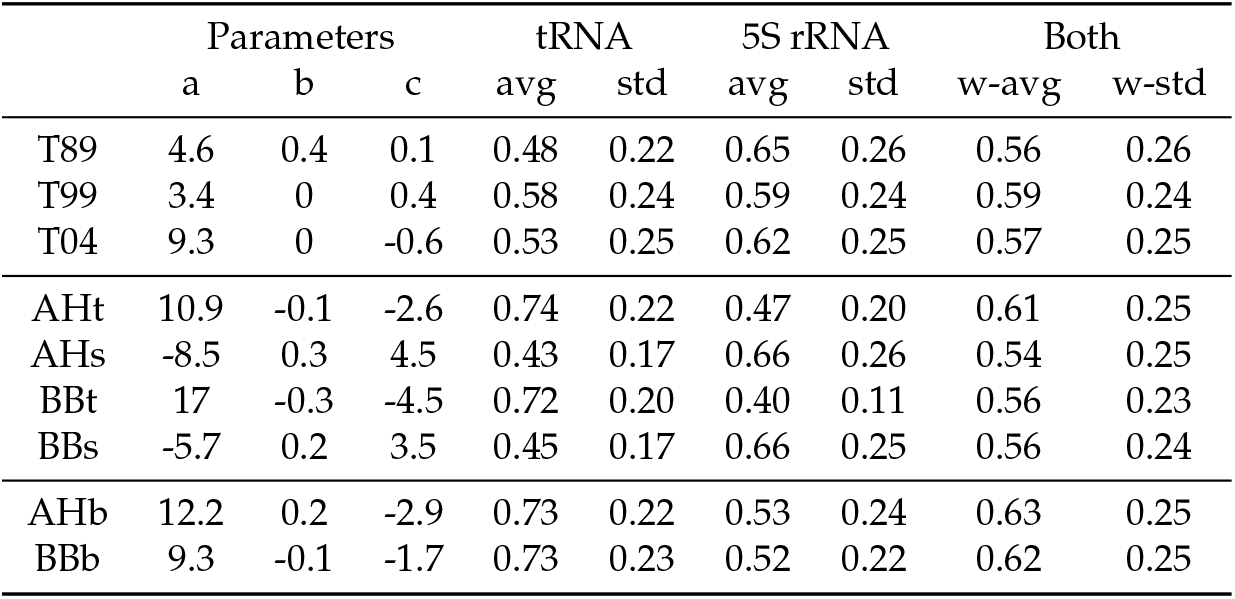
MFE prediction accuracies for testing data set from Mathews Lab (U Rochester) with 557 tRNA sequences and 1283 5S rRNA. Table 2 parameters repeated for completeness.

|  | Parameters |  |  | tRNA |  | 5S rRNA |  | Both |  |
| --- | --- | --- | --- | --- | --- | --- | --- | --- | --- |
|  | a | b | c | avg | std | avg | std | w-avg | w-std |
| T89 | 4.6 | 0.4 | 0.1 | 0.48 | 0.22 | 0.65 | 0.26 | 0.56 | 0.26 |
| T99 | 3.4 | 0 | 0.4 | 0.58 | 0.24 | 0.59 | 0.24 | 0.59 | 0.24 |
| T04 | 9.3 | 0 | -0.6 | 0.53 | 0.25 | 0.62 | 0.25 | 0.57 | 0.25 |
| AHt | 10.9 | -0.1 | -2.6 | 0.74 | 0.22 | 0.47 | 0.20 | 0.61 | 0.25 |
| AHs | -8.5 | 0.3 | 4.5 | 0.43 | 0.17 | 0.66 | 0.26 | 0.54 | 0.25 |
| BBt | 17 | -0.3 | -4.5 | 0.72 | 0.20 | 0.40 | 0.11 | 0.56 | 0.23 |
| BBs | -5.7 | 0.2 | 3.5 | 0.45 | 0.17 | 0.66 | 0.25 | 0.56 | 0.24 |
| AHb | 12.2 | 0.2 | -2.9 | 0.73 | 0.22 | 0.53 | 0.24 | 0.63 | 0.25 |
| BBb | 9.3 | -0.1 | -1.7 | 0.73 | 0.23 | 0.52 | 0.22 | 0.62 | 0.25 |

Other than this lower accuracy for 5S rRNA, several patterns observed in Table 2 also hold in Table 3. First, parameters which were trained on one family are markedly less accurate on the other one. When appropriately combined to weigh each family equally, this yields overall accuracies comparable to the Turner values. However, parameters chosen to raise the overall accuracy can achieve a statistically significant improvement over T99. Finally, branch-and-bound and ad hoc accuracies are remarkably similar over the families on which they were trained.

### 3.4. Sensitivity to parameter precision

As discussed in Section 2, the accuracy computations here focused on branching parameters specified to one decimal precision. We note that the “exact” parameters computed, that is ones specified as a rational number to the maximum precision allowable by the size of an integer in the computer algebra system used, always gave higher accuracy on the training (sub)set used. The largest difference seen was less that 0.035, which is not a statistically significant difference over data sets of this size with this amount of variance in the accuracy means.

## 4. Discussion

Previous results [17] demonstrated that it was possible to achieve a statistically significant improvement in MFE prediction accuracy by altering the three NNTM parameters which govern the entropic cost of loop branching. This was shown on a set of 50 tRNA sequences, on another of 50 5S rRNA, and on the full training set of both families combined. However, the extent of the possible improvement was unknown, although a lower bound was given by the ad hoc parameters identified — listed as AHt, AHs, and AHb respectively in Table 2. The “max” upper bound, known not to be attainable, was provided by the average maximimum accuracy (over any combination of parameters) for each individiual training sequence. Hence, the open questions was to establish the maximum *simultaneous* improvement over these training sets.

Here we provide a branch-and-bound algorithm which takes as input a set of RNA branching polytopes [23] and finds the parameters with the best possible accuracy over the entire set. We describe implementation details needed to insure that the basic algorithm runs efficiently enough to be useful in practice, and give results on our original training set as well as a much larger testing set available from the Mathews Lab (U Rochester) with 557 tRNA and 1283 5S rRNA. The differences in MFE accuracy under the standard T99 parameters between the training and testings sequences are not significant, and we find that the general trends observed in the training data are borne out by the testing results.

First, and most surprising, we find that the branch-and-bound parameters do not improve on the ad hoc ones in any significant way. Hence, we now know that the best possible MFE prediction accuracy for the tRNA training sequences is 0.75 on average and 0.73 for 5S rRNA. The testing data achieves a comparable accuracy for tRNA, although the ad hoc AHt parameters are actually slightly better than the branch-and-bound BBt. The AHs and BBs accuracies for the 5S rRNA testing sequences are equivalent, if lower that the training ones (but not by a statistically significant amount).

Overall, the average MFE prediction accuracy for the 5S rRNA testing sequences is consistently lower, by ∼ 5%, than the training accuracy for all parameter combinations considered. It is not obvious why this should be the case, since the GC content is equivalent and the length and number of native base pairs were actually lower on average for the testing sequences. In the future, it may be worthwhile to investigate what other sequence and/or structural characteristics might correlate with this training versus testing trend for 5S rRNA.

Returning to the accuracy improvement question, the branch-and-bound results for each family establish a much more realistic upper bound than the previous “max” values from Table 2. However, we also know that achieving this level of accuracy simultaneously across the two families is not possible. The testing data reinforces the point that parameters which are optimized only for one family perform much less well for the other. For the family-specific parameters, i.e. AHt, AHs, BBt, and BBs, this yields combined accuracies over both families which are on par with the Turner parameters for both training and testing data. In the case of the AHt paramaters, the improvement over T99 for the full testing set is statistically significant (*p* = 0.0187) but not for the training set (*p* = 0.1395).

In contrast, when the parameters are chosen to maximize the accuracy of both families, a statistically significant improvement over T99 — which has the highest combined accuracy over the three Turner parameter sets — is achieved for both testing and training data. As described for the training results, there are multiple different parameter combinations that achieve essentially the same accuracy over both families. This conclusion holds true for the testing data as well. Such stability in the maximum combined accuracy strongly suggest that future studies should focus on “near optimal” parameter combinations. This is particularly appropriate when the parameters are commonly specified to 1 decimal precision.

In general, we find that the AHb and BBb parameters are better for tRNA than for 5S rRNA, relative to the family-specific parameters. It is interesting to note that the Turner parameters with the highest 5S rRNA accuracy are the earliest ones which had *b* = 0.4, whereas the T99 and T04 parameters both have *b* = 0. The parameters trained only on 5S rRNA which produce the highest accuracy on that family, i.e. AHs and BBs, both had *b >* 0 while the opposite was true for tRNA. However, it was possible to achieve the same accuracy (to two decimal places) over the entire 100 sequences training set with either *b* = 0.2 or *b* = −0.1, and essentially the same for the 1840 sequence testing set.

Over all the new parameter combinations considered, we found a small range of *b* values, roughly centered around 0. Furthermore, previous results [17] had found that the thermodynamic optimization was most sensitive in the *b* direction. In future work, we expect to specialize the *b* value in our parametric analysis to better focus on the *a* and *c* trade-offs. We have preliminary results which indicate that this is not too detrimental to the overall accuracy, and this reduction in complexity will certainly improve the time and memory needed to compute the branching polytopes.

Recall that *a* weighs the number of multiloops while *c* scores the total number of branching helices. In terms of trade-offs between them, we note that the most recent Turner parameters have a significant increase in *a* and a *c <* 0 for the first time. All six new parameter combinations have opposite sign for *a* and *c* suggesting that there may be an important reward/penalty balance to be achieved. Not only did the 5S-specific parameters have *b >* 0 but they also both have *a <* 0 and *c >* 0, whereas the tRNA-specific had the exact opposite signs. Although the same accuracy could be achieved over the whole training data set with *b* either possitive or negative, there were no “near optimal” parameter combinations found which had *a <* 0 and *c >* 0. Hence, it seems that there may be a greater range of (*a, c*) combinations which produce an acceptable 5S rRNA accuracy than there is for tRNA. In fact, BBb is quite close to T04, with a difference of (0, − 0.1, − 1.1). This small change produces a considerable improvement in tRNA accuracy, with a smaller corresponding decrease for 5S rRNA.

In conjunction, these results suggest that there may be a relatively large set of branching parameters which yield equivalent prediction accuracies that improve over the current ones. To make progress on identifying the scope of these parameters, we confirm that the current empirical strategy of focusing on the trade-off between the *a* and *c* parameters is well-substantiated by this analysis. Moving forward, the goal is to expand the training set to include other RNA families, but this requires new algorithmic approaches to computing the branching polytopes. To date, the longest sequence attempted (an RNase P of length 354 nt) took more than 2 months. However, focusing on the *a,c* trade-offs only should reduce the complexity substantially, yielding further insights into the advances possible and limitation faced when improving RNA branching predictions.

## Author Contributions

Conceptualization, S.P. and C.H.; methodology, S.P., C.W., and C.H..; software, S.P., C.W., and M.C.; formal analysis, C.H.; investigation, S.P., C.W., and M.C.; writing— original draft preparation, C.W., and C.H.; writing—review and editing, S.P., and C.H.; supervision, S.P. and C.H.; funding acquisition, S.P. and C.H.. All authors have read and agreed to the published version of the manuscript.

## Data Availability Statement

The code developed and the data used in this study are openly available at https://github.com/spoznan/branching_polytopes_branch_and_bound.

## Abbreviations

The following abbreviations are used in this manuscript:

MFE: Minimum Free Energy
NNTM: Nearest Neighbor Thermodynamics Model

## Footnotes

1 After this step, for *L* = 0.74, the average number of regions to be considered for each of tRNA sequence is 12 and the total running time for tRNA on a machine with Intel^®^ Core™ i9-9900K, 32 GB SDRAM, is 2.5hrs. For *L* = 0.76, the average number of regions to be considered for each tRNA sequence is 9 and the total running time is 1.5hrs.

